# Molecular epidemiology and whole genome sequencing analysis of clinical *Mycobacterium bovis* from Ghana

**DOI:** 10.1101/489203

**Authors:** Isaac Darko Otchere, Andries J. van Tonder, Adwoa Asante-Poku, Leonor Sánchez-Busó, Mireia Coscollá, Stephen Osei-Wusu, Prince Asare, Samuel Yaw Aboagye, Samuel Acquah Ekuban, Abdallah Iddrisu Yahayah, Audry Forson, Akosua Baddoo, Clement Laryea, Julian Parkhill, Simon R Harris, Sebastien Gagneux, Dorothy Yeboah-Manu

## Abstract

**Background:** Bovine tuberculosis (bTB) caused by *Mycobacterium bovis* is a re-emerging problem in both livestock and humans. The association of some *M. bovis* strains with hyper-virulence, MDR-TB and disseminated disease makes it imperative to understand the biology of the pathogen.

**Methods:** *Mycobacterium bovis* (15) among 1755 *M. tuberculosis* complex (MTBC) isolated between 2012 and 2014 were characterized and analyzed for associated patient demography and other risk factors. Five of the *M. bovis* were whole-genome sequenced and comparatively analyzed against a global collection of published *M. bovis* genomes.

**Results:** *Mycobacterium bovis* was isolated from 3/560(0.5%) females and 12/1195(1.0%) males with pulmonary TB. The average age of *M. bovis* infected cases was 46.8 years (7-72years). TB patients from the Northern region of Ghana (1.9%;4/212) had a higher rate of infection with *M. bovis* (OR=2.7,p=0.0968) compared to those from the Greater Accra region (0.7%;11/1543). Among TB patients with available HIV status, the odds of isolating *M. bovis* from HIV patients (2/119) was 3.3 higher relative to non-HIV patients (4/774). Direct contact with livestock or their unpasteurized products was significantly associated with bTB (p<0.0001,OR=124.4,95% CI=30.1-508.3). Two (13.3%) of the *M. bovis* isolates were INH resistant due to the S315T mutation in *katG* whereas one (6.7%) was RIF resistant with Q432P and I1491S mutations in *rpoB*. *M. bovis* from Ghana resolved as mono-phyletic branch among mostly *M. bovis* from Africa irrespective of the host and were closest to the root of the global *M. bovis* phylogeny. *M. bovis*-specific amino acid mutations were detected among MTBC core genes such as *mce1A, mmpL1, pks6, phoT, pstB, glgP and Rv2955c*. Additional mutations P6T in *chaA*, G187E in *mgtC*, T35A in *Rv1979c*, S387A in *narK1*, L400F in *fas* and A563T in *eccA1* were restricted to the 5 clinical *M. bovis* from Ghana.

**Conclusion:** Our data indicate potential zoonotic transmission of bTB in Ghana and hence calls for intensified public education on bTB, especially among risk groups.

## Introduction

Among the *Mycobacterium tuberculosis* complex (MTBC), *Mycobacterium bovis* is the main causative agent of TB in cattle and sheep, albeit with the widest host range among other mammals including wildlife and humans [1]. *M. bovis* associated TB is a re-emerging global problem affecting both livestock and humans alike. The World Health Organization reported 147,000 new Bovine TB (bTB)) cases and 12,500 deaths among humans in 2016 [2]. Despite the low incidence of *M. bovis* associated TB (~2% globally), the mortality rate is high, especially among children and HIV co-infected patients [1,3,4]. Human-to-human transmission of *M. bovis* is mostly rare [5], thus human bTB is considered a zoonotic chronic disease characterized by lung infections and their draining lymph nodes as granulomatous necrotizing inflammatory disease [6,7]. Nevertheless, bTB among immunocompromised people and children are mostly extrapulmonary or disseminated affecting other organs other than the lungs and their draining lymph nodes. bTB in humans is mostly transmitted via the alimentary canal by the [4] consumption of unpasteurized dairy products from infected cattle [3,8,9] and or inhalation of aerosolised bacilli via direct contact with infected cattle and/or their carcasses [5]. However, a lack of knowledge or simply negligence of the dangers associated with being in close contact with livestock or wildlife and their unpasteurized products is apparent among some individuals who are constantly in direct contact with animals [10]. In addition, there is a growing association of *M. bovis* related TB cases with treatment failure due to intrinsic resistance to some commonly used anti-tuberculosis drugs [11].

Even though *M. bovis*, being a member of the MTBC, is genetically homogenous compared to other bacteria [12], molecular epidemiology of *M. bovis* infections in Great Britain has shown that they exhibit polymorphic metabolic profiles, such as differential rates of incorporation of propionate into membrane lipid components among different genotypes [13] as well as differential expression of some essential genes and accumulation of single nucleotide polymorphisms (SNPs) which could have functional implications [14].

About 85% of herds and 82% of humans live in close proximity in sub-Saharan Africa (SSA) in both rural and urban settings, driving the wide distribution of bTB compared to other global settings [15,16]. This is compounded by the inadequate sanitation practices such as the habit of sharing drinking water with beasts and consumption of non-pasteurized milk and dairy products [17–19] (. Despite the economic and public health importance of bTB, little knowledge exists on the epidemiology and biology of *M. bovis* in relation to the human adapted MTBC (hMTBC) lineages spanning *M. tuberculosis sensu stricto* (*Mtbss*) and *M. africanum* (*Maf*) [20,21]. However, such information is critical for development of effective control tools for bTB.

We determined the prevalence of bTB among pulmonary TB patients passively reporting to selected TB diagnostic/treatment facilities in Ghana, determined potential risk factors associated with bTB in Ghana and explored genomic similarities and differences among *M. bovis* strains from around the globe, irrespective of the host, using whole genome sequencing.

## Materials and Methods

### Ethical Statement and Participant Enrolment

The Institutional Review Board (IRB) of the Noguchi Memorial Institute for Medical Research (NMIMR) with Federal Wide Assurance number FWA00001824 reviewed this study and its protocols and accordingly gave ethical clearance in support of the work.

### Mycobacterial Isolation, Drug Resistance Profiling and Genotyping

Smear-positive sputum samples from the selected health centers in the Northern and Greater Accra regions of Ghana were decontaminated and inoculated on 2 pairs of Lowenstein Jensen (LJ) slants; one pair supplemented with 0.4% sodium pyruvate (to enhance growth of *M. bovis* and *M. africanum* (*Maf*)) the other with glycerol (for enhanced growth of *M. tuberculosis sensu stricto* (*Mtbss*) and incubated as previously described [22]. MTBC cells growing in confluence were harvested and heat inactivated at 95 °C for 60 min in nuclease-free water. After heat inactivation, chromosomal DNA was extracted using previously described protocol [23]. The isolates were confirmed as MTBC by PCR amplification of IS*6110* and spoligotyping was carried out for lineage classification [24]. Isolates classified as *M. bovis* were confirmed with a large sequence polymorphism (LSP) assay using PCR detection of deleted regions of difference RD9, RD4 and RD12 [25]. Drug susceptibility testing against isoniazid (INH) and rifampicin (RIF) was carried out using the micro-plate alamar blue assay [23,26].

### Whole Genome Sequencing and Phylogenetic Analysis

Whole genome sequencing of 5 candidate *M. bovis* isolates was carried out as previously described [27]. The 5 genomes (ERR502499; ERR502526; ERR502529; ERR502538; ERR1203064) were added to a collection of 767 previously published clinical and veterinary *M. bovis* genomes (supplementary data S1) from around the world for analysis. Sequence reads were mapped to the *Mycobacterium bovis* AF2122/97 reference genome (NC0002945) using BWA (minimum and maximum insert sizes of 50 and 1000 respectively) [28]. Single nucleotide polymorphisms (SNPs) were called using SAMtools mpileup and BCFtools (minimum base call quality of 50 and minimum root squared mapping quality of 30) as previously described [28,29]. Variant sites in the alignment were extracted using snp-sites [30] and a maximum likelihood phylogenetic tree was constructed using FastTree2 [31] (nucleotide general time-reversible tree). The resulting tree was annotated and rooted using iTOL [32]

### Comparative Mutational Analysis of Selected MTBC Core-Genes

Coordinates of 147 MTBC core genes (Supplementary Table S2) previously reported to harbour amino acid mutations with phenotypic consequence on virulence and fitness of some laboratory strains of the MTBC [33–40] were compiled from the Tuberculist database [41]. Using the fasta file of H37Rv as reference, the paired end reads of the 5 Ghanaian *M. bovis* genomes, 257 *M. africanum* [27] and global collection of 20 MTBC genomes [42] were screened for mutations within the compiled 147 core genes using ARIBA with default settings [43]. Amino acid mutations found to be present only among the 5 Ghanaian *M. bovis* genomes were suspected to be *M. bovis* specific. To confirm whether these mutations were widespread in *M. bovis*, the global collection of 767 clinical and veterinary *M. bovis* genomes (Supplementary data S1) was screened for these specific mutations using ARIBA as described above. We further classified these amino acid mutations as *M. bovis*-specific if they were found in 100% of genomes in the global collection or core *M. bovis* mutations if found in at least 99% of genomes.

### Statistical Analysis

Where applicable, chi-square and Fisher’s exact tests were used to establish statistical significance. *P-values* less than 0.05 were considered statistically significant with 95% confidence.

## Results

### Demography and Biological Associations of TB Patients infected with *M. bovis*

A total of 1755 MTBC isolates were obtained from 2074 smear positive TB patients (84.6% isolation rate). Among the patients from whom a MTBC was isolated, 212 (12.1%) were from the Northern region and 1543 (87.9%) from the Greater Accra region as previously described [27]. Fifteen (0.9%) of the isolates were genotyped as *M. bovis* whereas the remaining 1740 (99.1%) were members of the hMTBC (*Mtbss* and *Maf*). The average age of patients infected with *M. bovis* was 46.8 years (7 to 72 years) of which 12/1195 (1.0%) were from males compared to 3/560 (0.5%) from females (p = 0.412, OR = 1.9). Four (1.9%) of the isolates from the Northern region (n = 212) were *M. bovis* compared to 11/1543 (0.7%) from the Greater Accra region (p = 0.0968, OR = 2.7). Among the patients with known HIV status (893; 50.3%), 119 (13.3%) were HIV-positive compared to 774 (86.7%) HIV-negative. The incidence of bTB among HIV and non-HIV TB patients was 1.7% (2/119) and 0.5% (4/774) respectively with higher odds of isolating *M. bovis* from HIV patients relative to non-HIV TB patients (OR = 3.3). Six TB patients including 1 herdsman, 1 herds owner and 4 butchers representing 40% of 15 patients with history of direct contact with livestock were infected with *M. bovis*. This is significantly higher compared to 0.5% (9/1740) of *M. bovis* infected TB patients without such history (p < 0.0001, OR = 124.4, 95% CI = 30.1-508.3)

### Drug Resistance Profile of *M. bovis* Isolates

Most of the *M. bovis* isolates (13) were susceptible to all the drugs tested except two isolates resistant to INH and one isolate resistant to RIF (Table 1). The two INH resistant isolates both had the S315T mutation in *katG* while the RIF resistant isolate had Q432P and I1491S mutations in *rpoB*.

**Table 1:** Sensitivity of the MTBC Isolates to INH and RIF.

| Drug | Total (1755) | hMTBC (1740) | *M. bovis* (15) | *P-value* | OR | 95%CI |
| --- | --- | --- | --- | --- | --- | --- |
| INH<sup>r</sup> | 133; 7.6% | 131;7.5% | 2;13.3% | 0.3163 | 1.9 | 0.2-8.5 |
| RIF<sup>r</sup> | 16; 0.9% | 15;0.9% | 1;6.7% | 0.1288 | 8.2 | 0.2-61.0 |
| MDR | 40 (2.3%) | 40;2.3% | 0;0.0% | - | - | - |
| ANY | 189 (10.8%) | 186;10.9% | 3;20.0% | 0.2139 | 2.1 | 0.4-7.8 |

NB: ANY: Total number of isolates resistant to at least one drug.

### Phylogenetic Distribution of Global Collection of *M. bovis*

The maximum likelihood phylogenetic tree of global collection of *M. bovis* spanning both clinical and veterinary isolates rooted on *Maf* L6 as an outgroup shows random distribution of both the clinical and veterinary *M. bovis* (Fig 1). The majority of the global collection of *M. bovis* analyzed were isolated from animals (predominately cattle). The *M. bovis* genomes of African origin (Ghana, Eritrea and South Africa) generally clustered together closest to the root of the phylogeny irrespective of the host. Nevertheless, there were few *M. bovis* from South Africa which were sporadically distributed far from the root of the tree. There were 2 major clusters of *M. bovis* from New Zealand and one major cluster each from the United Kingdom, Mexico and the United States of America. Interestingly, the 5 Ghanaian clinical *M. bovis* clustered together as a monophyletic branch among the African *M. bovis* group (Fig 1).

**Fig 1:**
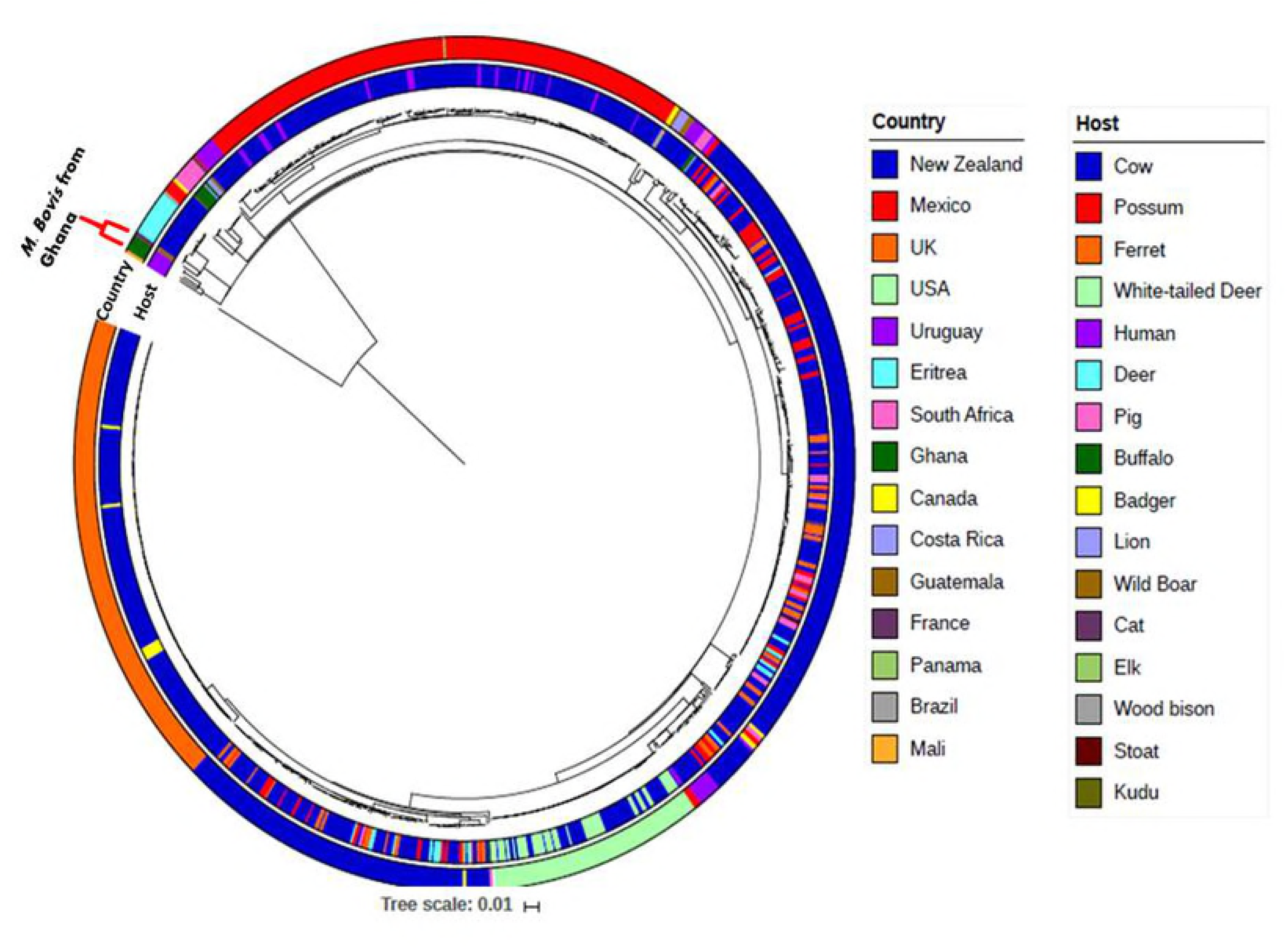
Phylogenetic tree of the Ghanaian clinical *M. bovis* amidst global collection of 767 published *M. bovis* genomes. The whole genome phylogeny of 767 publicly available M. bovis genomes together with 5 clinical M. bovis from Ghana rooted on M. africanum as an outgroup, shows the 5 Ghanaian clinical M. bovis genomes as a monophyletic group siting in a clade consisting mostly of other African M. bovis isolates basal to the rest of the dataset.

### *In silico* predicted *M. bovis*-Specific Amino Acid Mutations

We identified 41 *M. bovis* restricted amino acid mutations among 32 core-genes of the 5 clinical *M. bovis* from Ghana when compared to 257 *Maf* [27] and 20 global MTBC genomes [42] (Supplementary data S3). However, when we screened our global collection of 772 *M. bovis* genomes (including the 5 from Ghana), only 8 of the mutations were found in all genomes, 20 mutations in 99.22% to 99.87% of the genomes and 7 mutations in 95.85% to 98.97% of genomes. A further 6 mutations (P6T in *chaA*, G187E in *mgtC*, T35A in *Rv1979c*, S387A in *narK1*, L400F in *fas* and A563T in *eccA1*) were restricted to the 5 clinical *M. bovis* from Ghana (Fig 2; Supplementary Table S4; Supplementary Figure S5).

**Figure 2:**
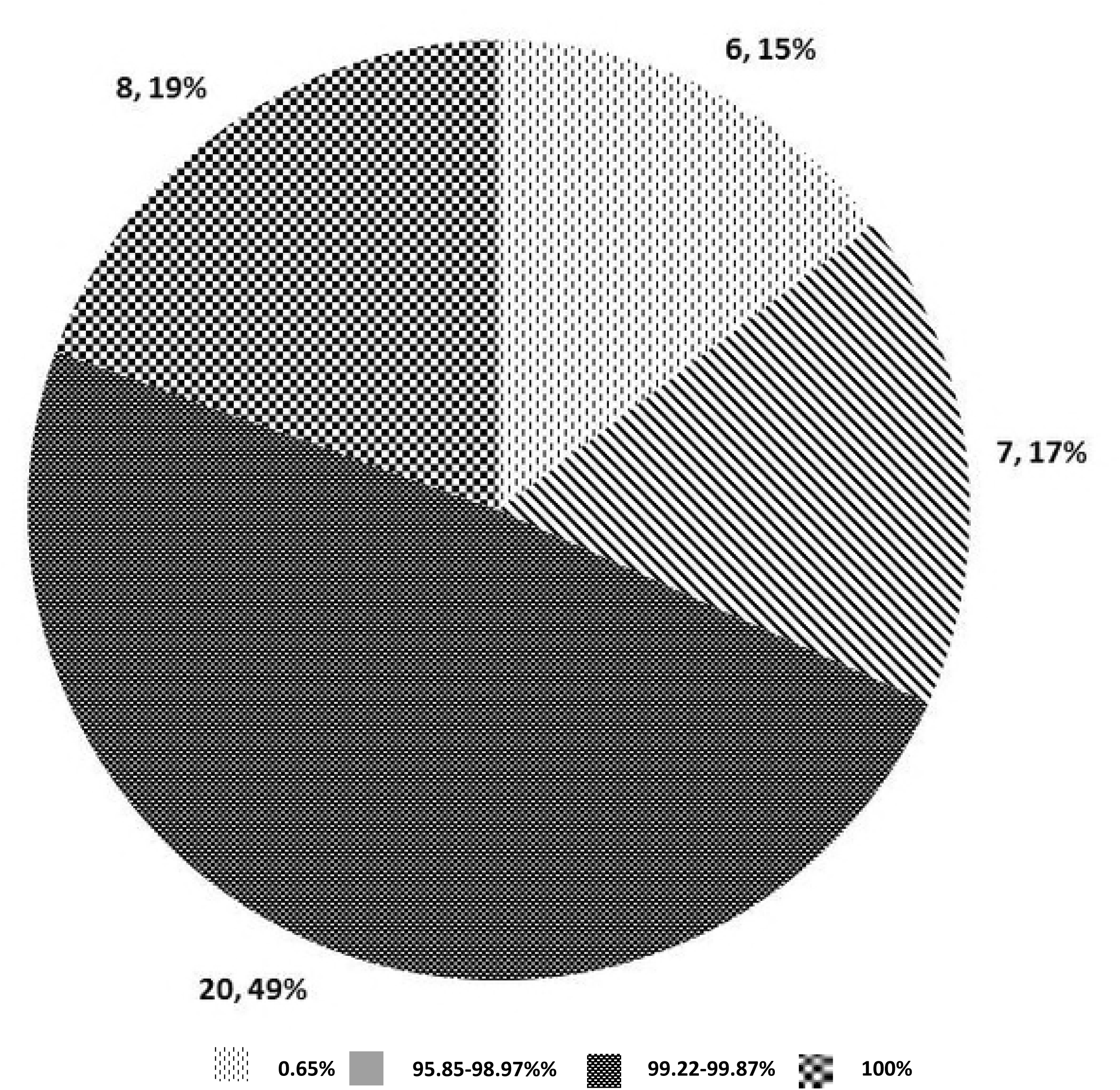
Distribution of selected core-gene amino acid mutations among *M. bovis*.

Among the 41 mutations identified uniquely among 32 core-genes *M. bovis*, 17 were among 15 essential genes associated with important physiological processes such as lipid metabolism, cell wall and cell processes, intermediate metabolism, and cellular respiration, virulence, detoxification and virulence as well as regulatory proteins (Table 2). These include *mce1A*, *phoT* and *eccA1* previously shown to be essential for the growth of *Mtbss* L4 strain H37Rv in primary murine macrophages [35]. In addition, mutations in other genes such as *pks6*, *pknD*, *pks4* and *glgP* have been shown to be associated with no production of phthiocerol dimycocerosates (PDIM) among mutant strains [36], attenuation in the central nervous system of BALB/c mice [40], no production of mycolic acid derivatives (mycolipanoic, mycolipenic and mycolipodienoic acids) among mutant strains [38] and *in vitro* slow growth [34].

**Table 2:**
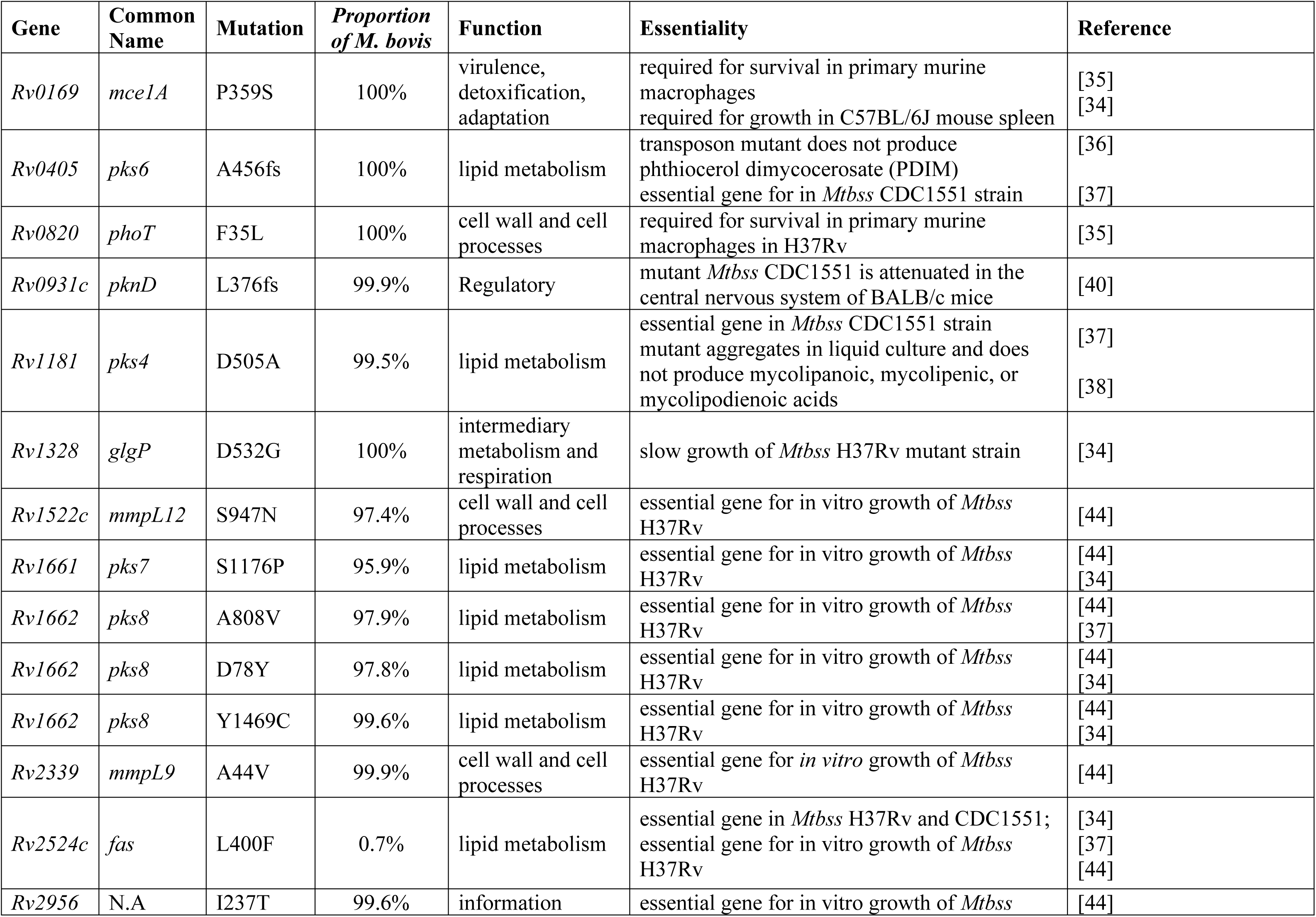
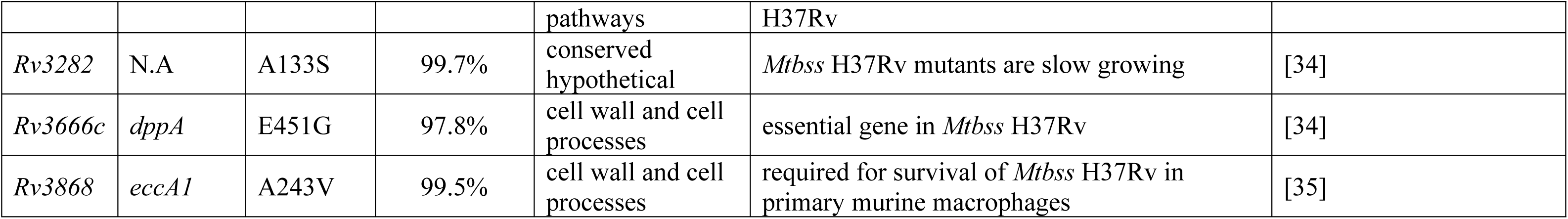
Description of *M. bovis*-Restricted Amino Acid Mutations among Essential Genes.

## Discussion

The global aim of reducing the impact of tuberculosis by the year 2030 cannot be achieved without considering the impact of zoonotic transmission and biology of *M. bovis*, the main causative agent of TB among cattle. The prevalence and incidence of bTB among humans is significantly lower across the globe compared to TB caused by the hMTBC [2]. Nevertheless, the association of bTB with compromised immunity and the innate resistance of *M. bovis* to pyrazinamide (PZA) (one of the four first line anti-TB drugs) underscore the need to adapt and implement TB control programs that encompass both bTB and TB caused by the hMTBC. Compared to other geographical regions, Africa has the highest burden of zoonotic transmission of bTB due to close contact of humans and animals (domestic and wild-life) as well as relatively poor hygienic practices [2,17,45–47]. We identified 15 *M. bovis* isolates among a total of 1755 MTBC isolated from pulmonary TB patients. Further molecular epidemiological analysis of these together with global collections of *M. bovis* and hMTBC showed (1) an association between close contact with livestock/animal carcasses and bTB infection in Ghana, (2) clustering of *M. bovis* of African origin close to the root of the global phylogeny and (3) the presence of *M. bovis*-specific amino acid mutations among both essential and non-essential core MTBC genes.

The finding of a significant association between bTB and close contact with animals (p < 0.0001) suggests zoonotic transmission and this calls for the implementation of preventive policies and strategies to reduce zoonotic transmission of TB among these high-risk groups [45]. This observation also calls for intensive education to create awareness of the disease about the risks of infection, the detection of infected animals/carcasses and prevention among farmers, butchers and the general population. Further emphasis should be placed on training butchers and animal handlers on the importance of adequate infection control measures, including the use of personal protective equipment (PPE) and the disposal of infected organs to avoid transmission of bTB among such personnel. An experienced butcher suffering from bTB in Australia gave an account of slaughtering many animals suspected of bTB and further cutting out the lungs for over 35 years without any proper precaution [48]. Also, some butchers in Nigeria, suffering from bTB, admitted eating visibly infected parts of the lung of cattle out of ignorance in order to convince customers to buy meat [49]. These instances highlight the importance of public education in the fight against bTB. This education should include veterinarians because there are instances of these professionals getting infected with bTB due to a lack of precautionary measures during execution of their work as was the case of a veterinary surgeon who suffered cutaneous bTB after performing several examinations without proper PPE [50].

Our observation also confirms the importance of the test and slaughter (TS) control strategy for bTB. In addition to pasteurisation of dairy products, bTB has been controlled in developed countries due to the successful implementation of the TS policy of all infected cattle and compensation of affected farmers by governments [51]. However, this has not been implemented in SSA due to the costs involved. Nevertheless, our findings call for a reconsideration of the TS strategy for bTB control in SSA and Governments must respond to this call.

We found the proportion of *M bovis* infected patients among participants from the Northern region (1.9%) of Ghana to be relatively higher (OR =2.7) compared to those from the Greater Accra region of Ghana (0.6%). The Northern region is home to over 70% of the national cattle population [52], confirming the observation that there is a relationship between close animal contact and bTB. Even though we found no clear association between the *M. bovis* isolates and drug resistance and HIV infection, the proportions were relatively higher than among the hMTBC. However, the lack of association may be because of the relatively limited number of *M. bovis* isolates thus further investigation using a larger number of isolates is required.

The global phylogeny of *M. bovis* clusters most of the *M. bovis* of African origin at the root of the tree (Fig 1) which might be an indication that they are closest to the progenitor of this successful member of the MTBC with the widest host range. However, the limited number of genomes from Africa does not allow inference of ancestry. With the exception of the five clinical *M. bovis* from Ghana which clustered as a monophyletic branch at the base of the tree, the random distribution of *M. bovis* irrespective of the speciation of the host underscores the wide host range of *M. bovis* and indicates that there is no specific host adaptation. However, the geographical distribution may suggest transmission of specific clones within certain geographical locations which agrees with earlier reports [53–55].

The identification and implications of *M. bovis*-specific amino acid mutations among genes such as *mce1A*, *phoT* and *eccA1* [35], *pks6* [36,38] as well as *glgP* [34] highlights the potential attenuated virulence of *M. bovis* relative to the hMTBC [56]. It would be interesting to test the effects of these mutations on fitness of mutants using *ex vivo* human cell lines or *in vivo* bovine models. In addition to the potential phenotypic implications of the identified mutations among essential genes, the 8 *M. bovis*-specific mutations could be utilized in developing either a rapid lateral flow diagnostic tool or a PCR-based tool specific for differential diagnosis of bTB among TB patients to advice an appropriate treatment regimen since *M. bovis* is innately resistant to pyrazinamide, a component of the DOTS regimen.

The scarcity of *M. bovis* genomes from African limited our ability to infer ancestry of the Ghanaian clinical isolates. Nevertheless, our data indicates a potential zoonotic transmission of bTB hence highlights the need for public education among people at risk. Moreover, the identified *M. bovis*-specific mutations could be utilized in the development of rapid diagnostic assays for differential diagnosis of bTB.

**List of Figure Legends**

**Acknowledgements**

**Author contributions**

## Competing financial interests

None declared

## Data availability

All the analyzed and/or generated data in this study are included in this article and its supplementary information files.

